# ToxCast chemical library screen identifies diethanolamine as an activator of Wnt signaling

**DOI:** 10.1101/2021.02.15.430319

**Authors:** Justin M. Wolter, Jessica A. Jimenez, Jason L. Stein, Mark J. Zylka

## Abstract

Numerous autism spectrum disorder (ASD) risk genes are associated with Wnt signaling, suggesting that brain development may be especially sensitive to genetic perturbation of this pathway. Additionally, valproic acid, which modulates Wnt signaling, increases risk for ASD when taken during pregnancy. We previously found that an autism-linked gain-of-function UBE3A^T485A^ mutant construct hyperactivated canonical Wnt signaling, providing a genetic means to elevate Wnt signaling above baseline levels. To identify environmental use chemicals that enhance or suppress Wnt signaling, we screened the ToxCast Phase I and II libraries in cells expressing this autism linked *UBE3A^T485^* gain-of-function mutant construct. Using structural comparisons, we identify classes of chemicals that stimulated Wnt signaling, including ethanolamines, as well as chemicals that inhibited Wnt signaling, such as agricultural pesticides, and synthetic hormone analogs. To prioritize chemicals for follow-up, we leveraged predicted human exposure data, and identified diethanolamine (DEA) as a chemical that both stimulates Wnt signaling in *UBE3A^T485A^*–transfected cells and has a high potential for prenatal exposure in humans. DEA also enhanced proliferation in two primary human neural progenitor cell lines. Overall, this study identifies chemicals with the potential for human exposure that influence Wnt signaling in human cells.

## Introduction

Large-scale exome sequencing studies of individuals with autism identified over 100 high-confidence ASD genes(1–3). Approximately 19% of these ASD genes are associated with the Wnt/ß-catenin signaling pathway, suggesting that alterations in Wnt signaling contribute to ASD pathogenesis(4–9). Members of the Wnt family are secreted signaling proteins that affect the development of nearly every area of the central nervous system(10). In the developing brain, Wnt establishes the anterior/posterior and dorsoventral axes, and instructs cell fate decisions by regulating the balance between differentiation and proliferation(11). Constitutive activation of Wnt signaling leads to hyperproliferation of neural progenitor cells and macrocephaly(12).

Non-genetic environmental factors also contribute to autism risk(6, 13, 14). Epidemiological studies link gestational exposure to agricultural pesticides with risk for ASD(15, 16). And, certain environmental-use chemicals can mimic transcriptional changes associated with ASD when applied to primary mouse neuron cultures(17, 18). The best characterized environmental risk factor for ASD is valproic acid (VPA), which is prescribed for epilepsy, bipolar depression, and migraine(19). Prenatal exposure to VPA increases the risk of congenital malformations(20), ASD(21, 22), and macrocephaly(23, 24). VPA activates Wnt signaling by targeting HDAC1(25). Furthermore, drugs approved by the FDA for treating behavioral symptoms of ASD (aripipazole, risperidone) can affect Wnt signaling(26, 27). These studies suggest that the developing nervous system may be highly sensitive to chemicals in the environment that modulate Wnt signaling.

Identifying environmental risk factors for neurodevelopmental disorders is a major challenge due to the lack of developmental neurotoxicological data on the vast majority of chemicals(28). To address this critical need, the EPA created the Tox21 program, which aims to provide platforms and methods to rapidly screen chemicals for potential adverse health effects(29). Here, we hypothesized that Wnt modulating chemicals will have enhanced effects in cells expressing an ASD-linked gene that, when overexpressed, stimulates Wnt signaling. To test this hypothesis, we screened the EPA Toxcast Phase I/II libraries using a Wnt sensitive luciferase reporter(30) in cells overexpressing UBE3A with an autism-linked T485A mutation (*UBE3A^T485A^*), a mutation that promotes Wnt signaling(31, 32).

## 2. Materials and Methods

### 2.1 Lentiviral infection of primary mouse cortical neurons

All lentivirus was produced in HEK293T cells using the third-generation packaging plasmids(33). Supernatant was collected, filtered using 0.45 μM filters, and frozen in single use aliquots.

Primary neuron cultures from E15.5 C57Bl/6 mouse embryos were prepared as previously described(18). Neurons were plated in 96 well plates at 20,000 cells per well. On day three, cells were infected with lentiviruses carrying BAR:luciferase and Tk:*Renilla* in a 5:1 ratio. Cells were incubated for five days, then treated with ToxCast chemicals and incubated for 48 hours. Cells were lysed and the lysate was used in dual luciferase assays using the Dual-Glo luciferase system (Promega), and measured on the GloMax Discover plate reader (Promega).

### 2.2 High-throughput Wnt screen of ToxCast phase I/II libraries

All liquid handling steps of HEK293T ToxCast Phase I/II screen were performed using the Tecan EVO liquid handling robot. These steps included cell plating, chemical library dilution and aliquots, cell dosing, transfections, and luciferase assays. Technical replicates for six control chemicals (three Wnt inhibitors and three Wnt activators) were spiked into random positions in each plate to ensure technical reproducibility and eliminate the risk of plate swaps. HEK293T cells were cultured in DMEM (Gibco) and 10% FBS in the absence of antibiotic in a humid incubator at 37°C with 5% (vol/vol) CO_2_. Cells were plated in white opaque 384 well plates at a density of 4,500 cells per well. 24 hours post plating cells were transfected with a β-catenin responsive luciferase reporter (BAR)(30), TK-*Renilla*, and pCIG2 *UBE3A^T485A^* using Fugene 6 (Promega). 4 hours post transfection cells were treated with chemical libraries. Cells were lysed 24 hours later, and luciferase assays were performed using the Dual-Glo luciferase system (Promega). All steps, including cell culture, treatments, lysis, and luciferase assays were performed in the same plate to minimize technical variation from handling artifacts. Four biological replicates (one well per chemical per concentration per day) were performed on different days to ensure reproducibility and reduce batch effects.

### 2.3 Screen analyses

The “Wnt luciferase ratio” was calculated by dividing the raw Firefly value by the raw *Renilla* luciferase value, and median centering within each plate. “Cell Health” was calculated using the raw *Renilla* value median centered within each plate. Biological replicates were averaged, and *P*-values were calculated using a two tailed T-test. To calculate the “Wnt Score” (Wnt activity with a penalty for toxicity) we calculated the mean log2 fold change of the Wnt luciferase ratio for each chemical, calculated the slope of the concentration-response curve for that chemical, and multiplied this by the mean of the Cell Health metric. The EPA spiked in replicate chemicals across plates to assess reproducibility, in addition to the six control chemicals we added. When a chemical was present in multiple plates we averaged the values for each metric.

### 2.4 HEK293T versus neuron toxicity comparison

RASL-seq assessed Toxcast Phase I chemical toxicity in primary mouse neuron cultures by spiking in control luciferase RNA in each well, and calculating the ratio of luciferase reads to total number of reads from neurons(18). We normalized this data by median centering and averaged the values for all concentrations of each chemical. We then compared the measure of cell health from primary neurons to the Cell Health Metric from this screen.

### 2.5 Structural chemical clustering

Chemical structuring was performed using ChemMineR(34). SMILE strings were converted to SDF files. Distance matrices were defined using atom-pair properties, and unsupervised hierarchic clustering was performed using R.

### 2.6 Estimated human exposure data

Estimated human exposure data was downloaded from(35). We compared the Wnt Score metric with the 95% confidence interval mg/kg/body weight/day for reproductive age females (defined as 16-49 years old), reasoning that this demographic is most representative of maternal, fetal, and neonatal exposure.

### 2.7 DEA in HEK293T cells

DEA was obtained from Sigma-Aldrich (#31589). The panel of luciferase reporter plasmids was a kind gift from the lab of Dr. Ben Major. Luciferase assays were performed as described above. The following day cells were transfected with either pCIG2:empty (eGFP with an IRES carrying empty sequence) or pCIG2 *UBE3A^T485A^*. Four hours later cells were treated with the indicated concentrations of DEA or vehicle (DMSO). Cells were incubated for 48 h and then total RNA was extracted using Trizol. cDNA was synthesized using SuperScript IV VILO with ezDNase (ThermoFisher). qPCR experiments were performed using SsoAdvanced Universal SYBR Green Supermix (NEB) on the Quantstudio5 (Applied Biosystems). Data was normalized to *EIF4A2* using the ∆∆Ct method. Two-tailed t-tests were used for comparison between vehicle conditions, and two-way ANOVA was used for concentration-response curves.

### 2.8 Primary human neural progenitor cell cultures

Human fetal brain tissue was obtained from the UCLA Gene and Cell Therapy Core following IRB regulations. Primary human (ph)NPCs were grown and differentiated as previously described(36, 37). Briefly, cells were thawed and plated in 10 cm plates with proliferation media (Neurobasal A supplemented with primocin, BIT9500, glutamax, heparin, EGF, FGF, LIF, PDGF) in a humid incubator at 37°C with 5% (vol/vol) CO_2_. Cells were mycoplasma tested and confirmed to be mycoplasma free (ATCC, Universal Mycoplasma Detection Kit). For experiments in Supporting Figure 1b cells were plated in 96 well plates and infected with lentivirus carrying BAR:luciferase and Tk:*Renilla* in a 5:1 ratio. Cells were incubated for 48 hours, then treated with the indicated chemicals. Cells were incubated for an additional 48 hours, then lysed and subjected to dual luciferase assays, as described above. For experiments in Figure 5 cells were plated in 96 well plates at a density of 12,500 cells per well. 24 hours later cells were treated with DEA, and incubated for 46 hours. We then performed a two-hour pulse with 10 uM EdU, then fixed the cells with 4% paraformaldehyde. Labelling was performed using the Click-iT EdU fluorescent labeling kit per manufacturer’s instructions (Thermo-Fisher Cat. C10337). DNA was labeled using FxCycle Far Red stain (Invitrogen, Cat# F10348). Cells were counted using the Attune NxT. Data was analyzed using the FlowJo software.

**Figure 5.**
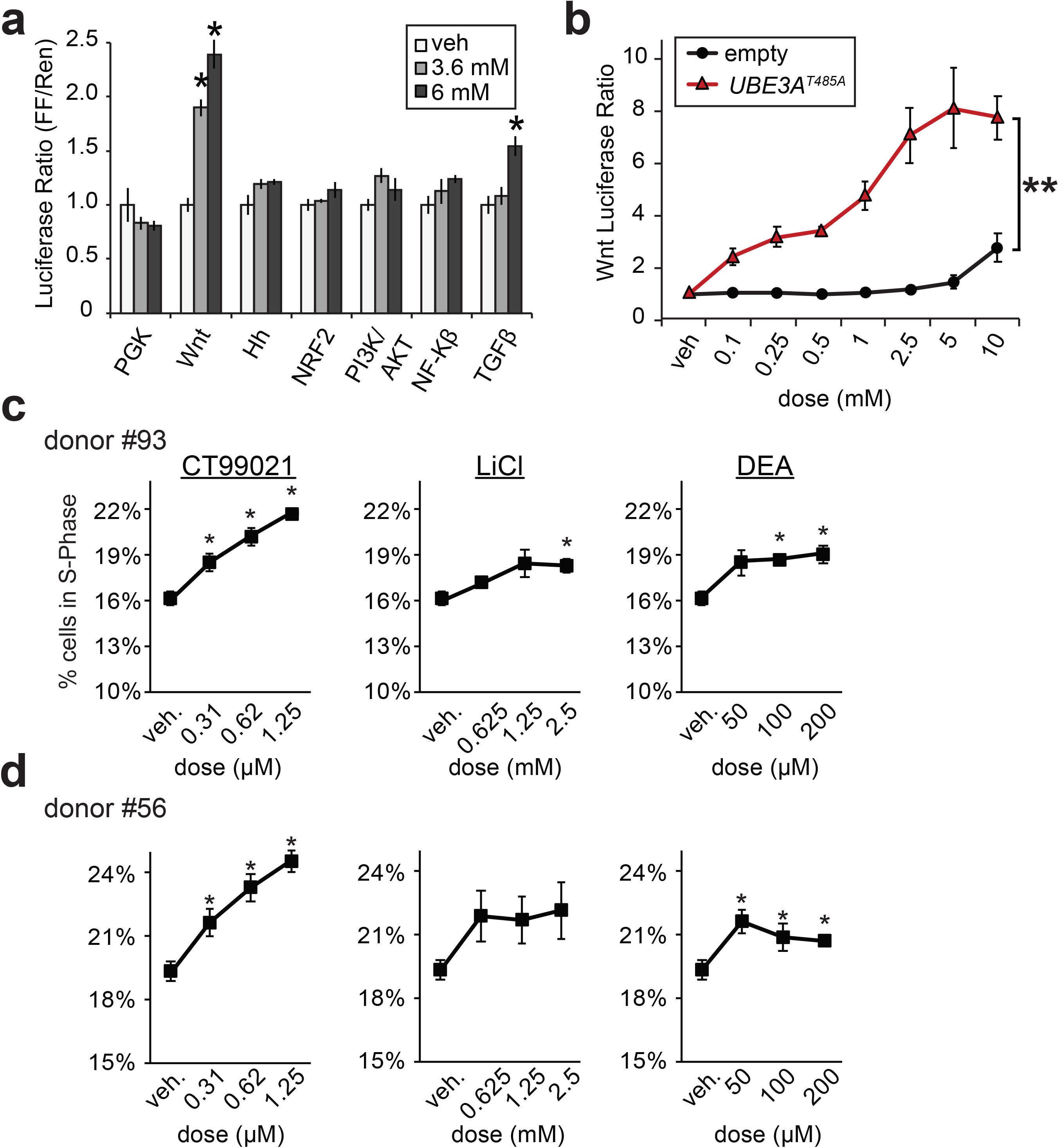
DEA activates Wnt signaling and proliferation. **a)** The effect of DEA on several luciferase reporters that measure developmental signaling pathways. Experiments done in the absence of *UBE3A^T485A^* overexpression. Tk:Renilla cotransfected for internal control. PGK (ubiquitous promoter, negative control), Hh (Hedgehog). Data normalized to vehicle for each reporter. T-test, * p<0.05, n = 4. **b)** Concentration-response curve of DEA on Wnt luciferase reporter in the presence of either empty plasmid, or *UBE3A^T485A^* overexpression. ANOVA, effect of genotype on Wnt response, ** p<0.01. **c,d)** Proliferation rates of Wnt control chemicals and DEA in two primary human neural progenitor cell lines. Cells treated for 46 hours with indicated chemical and concentration, followed by a two hour pulse with EdU. Cells analyzed by flow cytometry. T-test, * p<0.05, n = 4.

## 3. Results

### 3.1 High-throughput screen for environmental-use chemicals that modulate Wnt signaling

Given the evidence implicating Wnt signaling in ASD pathogenesis, we set out to test the EPA ToxCast Phase I and Phase II libraries(38) in cells transfected with an ASD linked *UBE3A^T485A^* mutant expression construct(31). Toxcast libraries contain chemicals with the potential for human exposure, including pesticides, plasticizers, perfluorinated chemicals, and “failed-pharma” compounds, which were donated by pharmaceutical companies due to toxicity in trials(38). We were blind to the identities of ToxCast Phase II chemicals during the screen, and were only unblinded after sharing the results of our screen with the EPA.

To quantify Wnt signaling, we used the β-catenin activated reporter (BAR) luciferase reporter, which contains 12 tandem binding sites for the TCF/LEF transcription factor(30). We cotransfected a *Renilla* luciferase reporter driven by the Thymidine Kinase (TK) promoter as an internal control to assess cell viability and toxicity. Overexpression of *UBE3A^T485A^* activates the Wnt reporter by inhibiting proteasome dependent degradation of β-catenin(32). To identify a representative cellular context in which to perform the screen, we tested known Wnt activators in primary mouse cortical neurons, primary human neural progenitor cells (phNPCs), and HEK293T cells (Supporting Figures S1a-c). Control chemicals included VPA(25), the GSK3β inhibitor CT99021(39) and lithium chloride(40). We found context specific effects, with LiCl not activating the Wnt reporter in primary mouse neurons (Supporting Figure S1a), and VPA not activating the Wnt reporter in phNPCs (Supporting Figure S1b). HEK293T were the only cells that demonstrated Wnt activation of all three chemicals, therefore we chose these cells to perform the screen (Supporting Figure S1c). Wnt inhibitors and activators received a positive Z-factor, a statistical measure of assay suitability for high-throughput screening(41) (Supporting Figure S1d).

Our two endpoints were Wnt luciferase ratio (BAR/*Renilla*, Figure 1a) and “cell health” (*Renilla* values, Figure 1b) (see Materials and Methods). We considered putative Wnt modulators as those with log2 fold change abs(log2 fold change) > 1 compared to vehicle, and p-value<0.05 (Figure 1a, Supporting Table S1). All control chemicals performed as expected (arrows, Figure 1a).

**Figure 1.**
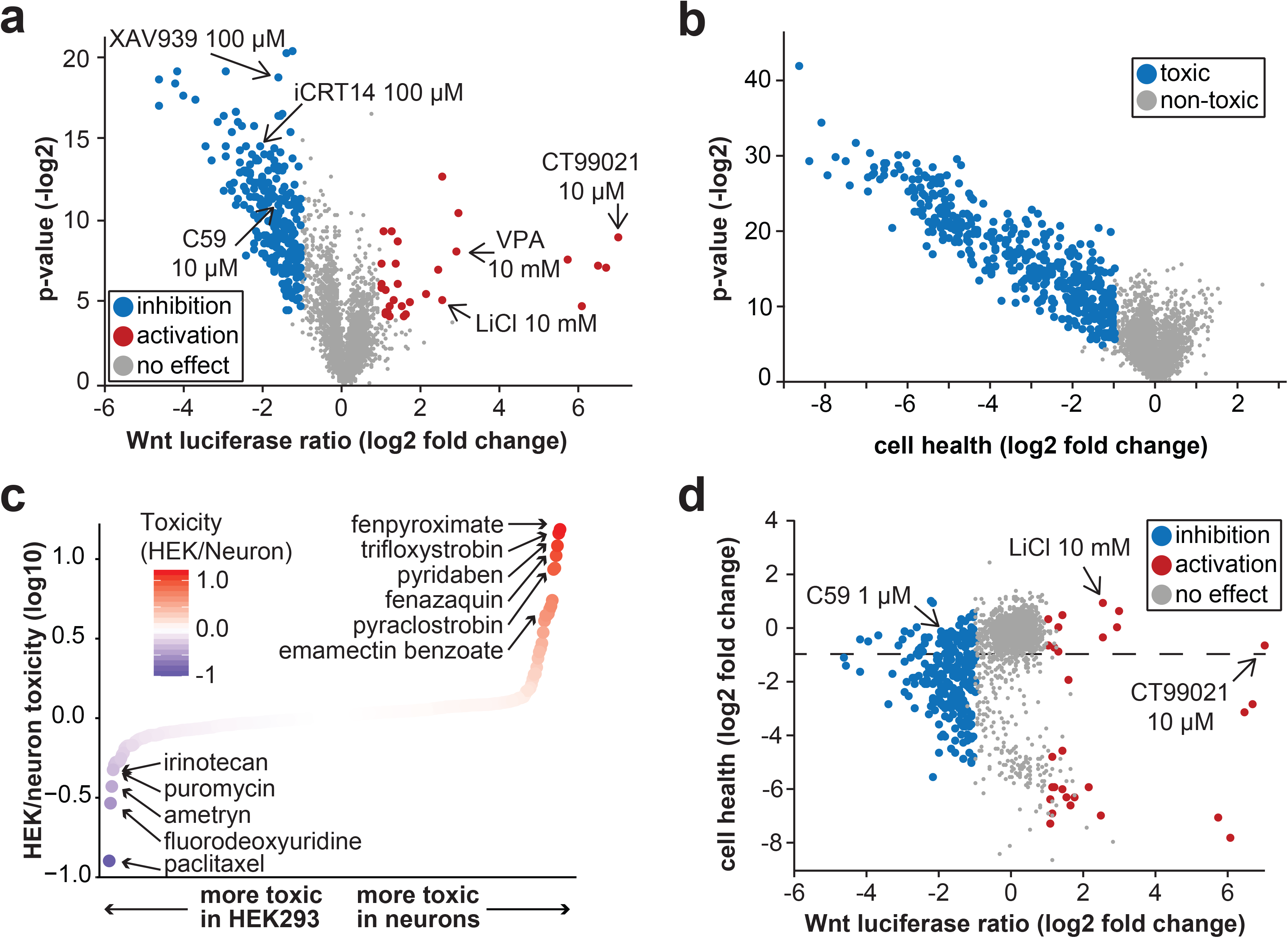
Screen to identify ToxCast chemicals that stimulate or inhibit Wnt signaling. **a)** ToxCast phase I/II chemicals screened against the Wnt luciferase reporter in HEK293T cells transfected with *UBE3A^T485A^* expression plasmid. Arrows mark chemicals that were used as positive controls. Each point is a single chemical at a single concentration. P-value represents unpaired T-test comparing each chemical with negative control vehicle wells in each plate. **b)** Cell health of ToxCast chemicals in HEK293T cells transfected with *UBE3A^T485A^* expression plasmid. Each point is a single chemical at a single concentration. Decrease in cell health score indicates toxicity. **c)** Comparison of ToxCast chemical toxicity in HEK293T cells transfected with *UBE3A^T485A^* and primary mouse neuron cultures. Toxicity was calculated as the slope of *Renilla* luciferase (internal control) signal across all concentrations of each chemical. **d)** Comparison of cell health and Wnt activation measures. Each point is a single chemical at a single concentration. Chemicals below the dashed line are those that have toxic effects.

### 3.2 Toxicity of ToxCast phase I/II chemicals

Many of the ToxCast chemicals exhibited concentration-dependent toxicity (log2 fold change < −1, and p-value<0.05, Supp. Table 1, Figure 1b). Previously, we tested the ToxCast Phase I library, which contains mostly pesticides(38), in primary mouse neuron cultures using RNA-seq as well as RASL-seq—a massively pooled transcriptomic assay(17, 18). In the RASL-seq experiments we also estimated neuronal toxicity by comparing total read counts per well to a luciferase mRNA spike in control. To identify chemicals with context specific toxicity, we compared the toxicity values in HEK293T cells (Figure 1b) with those in primary neurons (Figure 1c). The chemicals which were specifically toxic in HEK293T cells were mechanistically broad (Supporting Table S2), but typically exert anti-mitotic effects, such as the chemotherapeutics paclitaxel, fluorodeoxyuridine, and irinotecan(42, 43). Among these chemicals were also environmental use pesticides such as ametryn, the most widely used herbicide in sugarcane production and a frequent contaminant in aquatic environments(44, 45). In contrast to the broad mechanisms of toxicity in HEK293T cells, the chemicals that were most toxic to neurons were mitochondrial complex I and III inhibitors. These included fenpyroximate, trifloxystrobin, pyridaben, fenazaquin, and pyraclostrobin (Figure 1c, Supporting Table S2)(46, 47). This class of chemicals is functionally related to rotenone (Supporting Table S2), which is implicated in Parkinson’s disease(48, 49). Emamectin benzoate, a chemical that binds with high affinity to invertebrate GABA receptors(50, 51), was also selectively toxic in mouse neurons.

We next compared Wnt modulation with toxicity, and found that many chemicals that activated or inhibited the Wnt reporter were also toxic (Figure 1d). This is in contrast to our control chemicals which modulated Wnt without strong toxicity (Figure 1d, Supporting Figure 2a,b). Therefore, we generated a metric termed the “Wnt Score,” which reflects the potency of each drug across multiple concentrations with a penalty for toxicity (See Materials and Methods, Supporting Figure S2). All the control chemicals segregated to the top of this list (Figure 2a). To identify high-confidence non-toxic Wnt modulators, we filtered for those with p<0.05, log2 fold change greater than abs(log2 fold change) > 1, and Wnt score >0.4.

**Figure 2.**
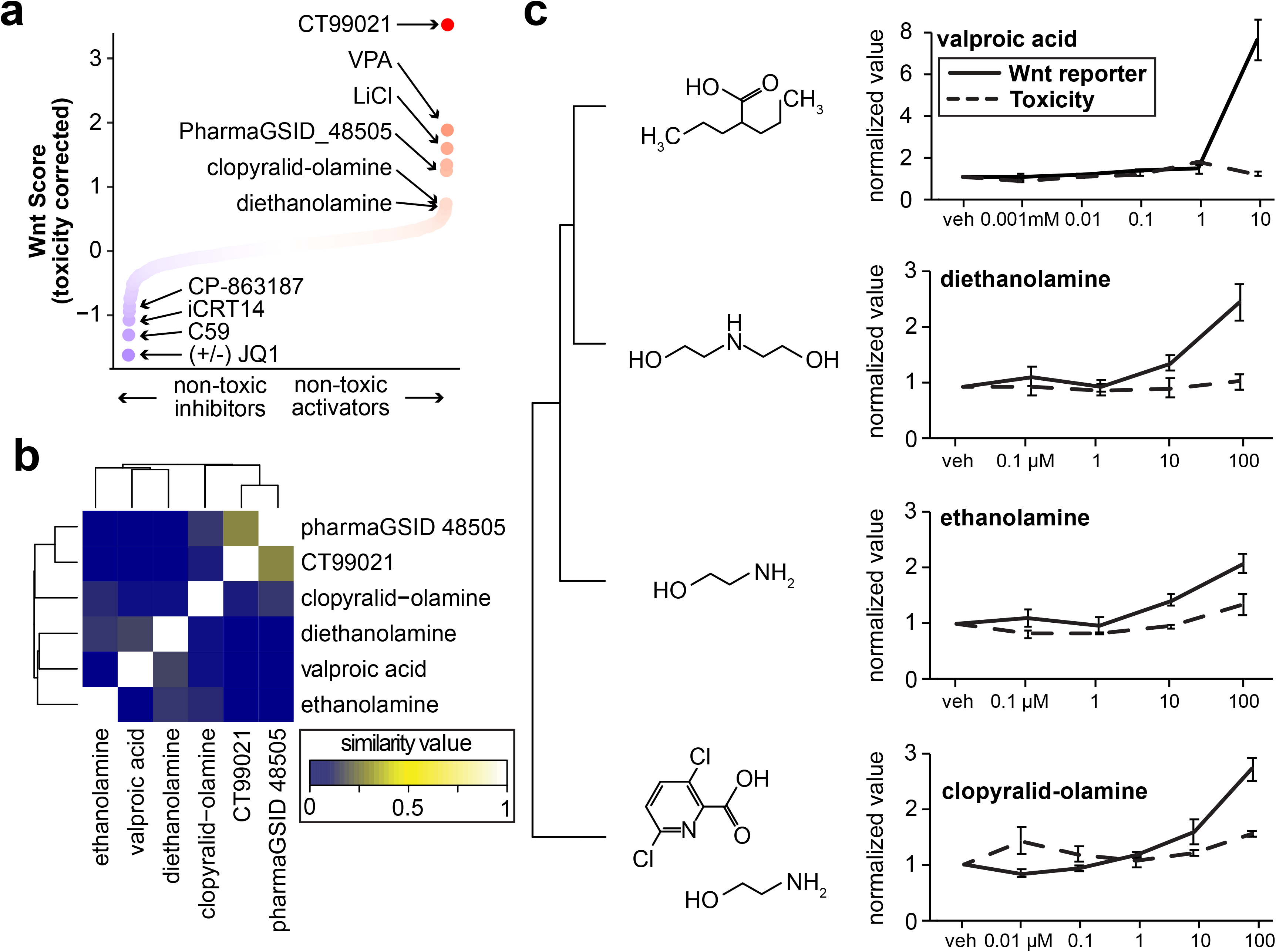
Non-toxic Wnt activators. **a)** Toxicity corrected Wnt luciferase ratio (Wnt Score), which combines all concentrations of each chemical and imparts a penalty for toxicity (mean luciferase ratio of all concentrations (log2 fold change), multiplied by the mean *Renilla* values for all concentrations). Positive control chemicals for both activation and inhibition rose to the top of this list. **b)** Comparison of chemical structures of non-toxic Wnt activators using SMILE strings and hierarchical clustering. **c)** Concentration-response curves for Wnt luciferase signal and toxicity scores for the ethanolamine cluster. Values normalized to vehicle.

### 3.3 Structural and functional comparisons of non-toxic Wnt modulators

Structural comparisons of chemical libraries can be used to group chemicals with similar structures to infer common functions and molecular targets. To characterize structural similarities in the ToxCast chemicals, we used SMILE strings to performed hierarchical clustering and multidimensional scaling(34). The most potent Wnt activator in the Toxcast library was pharmaGSID_48505, which has structural similarity with CT99021 (Figure 2b, Supporting Figure S3a,b). The similarity in effect size and structure between these two molecules suggests pharmaGSID_48505 targets GSK3β, but the enhanced toxicity suggests it is not as specific as CT99021 (Supporting Figs. 3a,b). The next cluster of Wnt activators contains several forms of ethanolamine (Figure 2b,c). Ethanolamines are bifunctional chemicals, containing a primary amine group and a primary ethanol group. Ethanolamine forms the head group of the phospholipid phosphatidylethanolamine, which is highly abundant in the inner leaflet of cell membranes(52), and comprises ~45% of all phospholipids in the brain(53). Both ethanolamine and diethanolamine (DEA) activated the Wnt reporter without toxic effects (Figure 2c), while triethanolamine had no effect (Supporting Figure S3c). DEA has marginal structural similarity to VPA (Figure 2b). Clopyralid-olamine, a mixture of clopyralid and ethanolamine, also activated the Wnt reporter (Figure 2c). However, clopyralid alone had no effect (Supporting Figure S3d), suggesting that ethanolamine in this mixture was responsible for activating the Wnt reporter.

Wnt inhibitors were substantially more numerous than activators, highlighting the benefit of screening the Wnt reporter in cells transfected with *UBE3A^T485A^*, which activates Wnt signaling (Figure 3a). Multiple agricultural pesticides inhibited the Wnt reporter, and these were structurally diverse (Figure 3a). These included the mitochondria complex I inhibitor tebufenpyrad (Figure 3b), and flufenacet, which inhibits synthesis of very long chain fatty acids (Figure 3c)(54, 55). Three inhibitors of p38 were also identified (Figure 3a), including CP-863187 which is a highly potent and selective p38 inhibitor (Figure 3d)(56). P38 regulates the canonical Wnt pathway through GSK3β(57), again highlighting GSK3β as a central regulatory node of the Wnt pathway. Four clusters resolved when comparing chemical similarity, including synthetic estrogens (Figure 3e), thyroid hormone analogs (Figure 3f), glucocorticoid and steroid hormones (Figure 3g), and agricultural fungicides (Figure 3h). The crosstalk between these hormone signaling pathways and Wnt signaling is well established(58–62). These results raise the possibility that exposure to multiple chemicals with structural and functional similarity might have additive effects by acting through the same molecular pathways.

**Figure 3.**
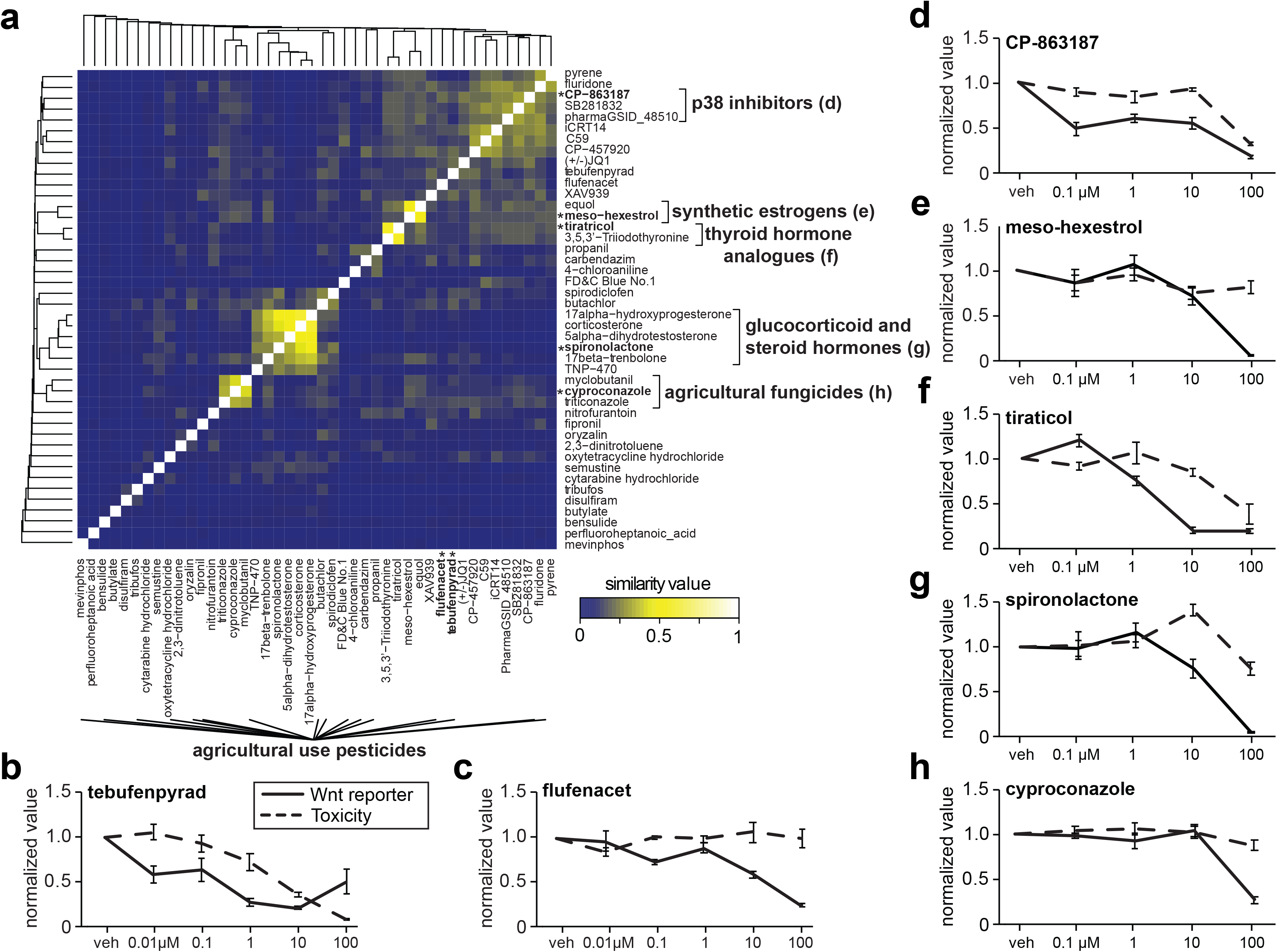
Non-toxic Wnt inhibitors. **a)** Comparison of chemical structures of non-toxic Wnt inhibitors using SMILE strings and hierarchical clustering. Representative chemicals displayed in **b-h** marked by asterisks. **b-h)** Concentration-response curves for Wnt luciferase signal and toxicity scores for representative chemicals of each class.

### 3.4 Prioritizing chemicals using predicted human exposure data

Humans are exposed to thousands of environmental-use chemicals, yet exposure data is not available for the majority of these chemicals(63). Instead, exposure estimates can be generated using various parameters, including urine biomonitoring of representative chemicals, chemical use classes, and production volume(35). We used these estimates to prioritize chemicals for more detailed validation experiments (Figure 4). We focused on exposure (mg/kg/body weight/day) predictions for reproductive age (16–49) females, reasoning that this age group best represents in utero exposure estimates (Figure 4, Supporting Table S3). The inhibitor with the highest relative exposure predictions was FD&C Blue No.1 (Figure 4). This dye has been approved for use in foods since the early 1900’s, and is considered safe and non-toxic by the FDA. It is deep blue in color, which visibly altered the color of cell media, which could interfere with the sensitivity of the luciferase assay. For these reasons we did not pursue this chemical for further experimental validation.

**Figure 4.**
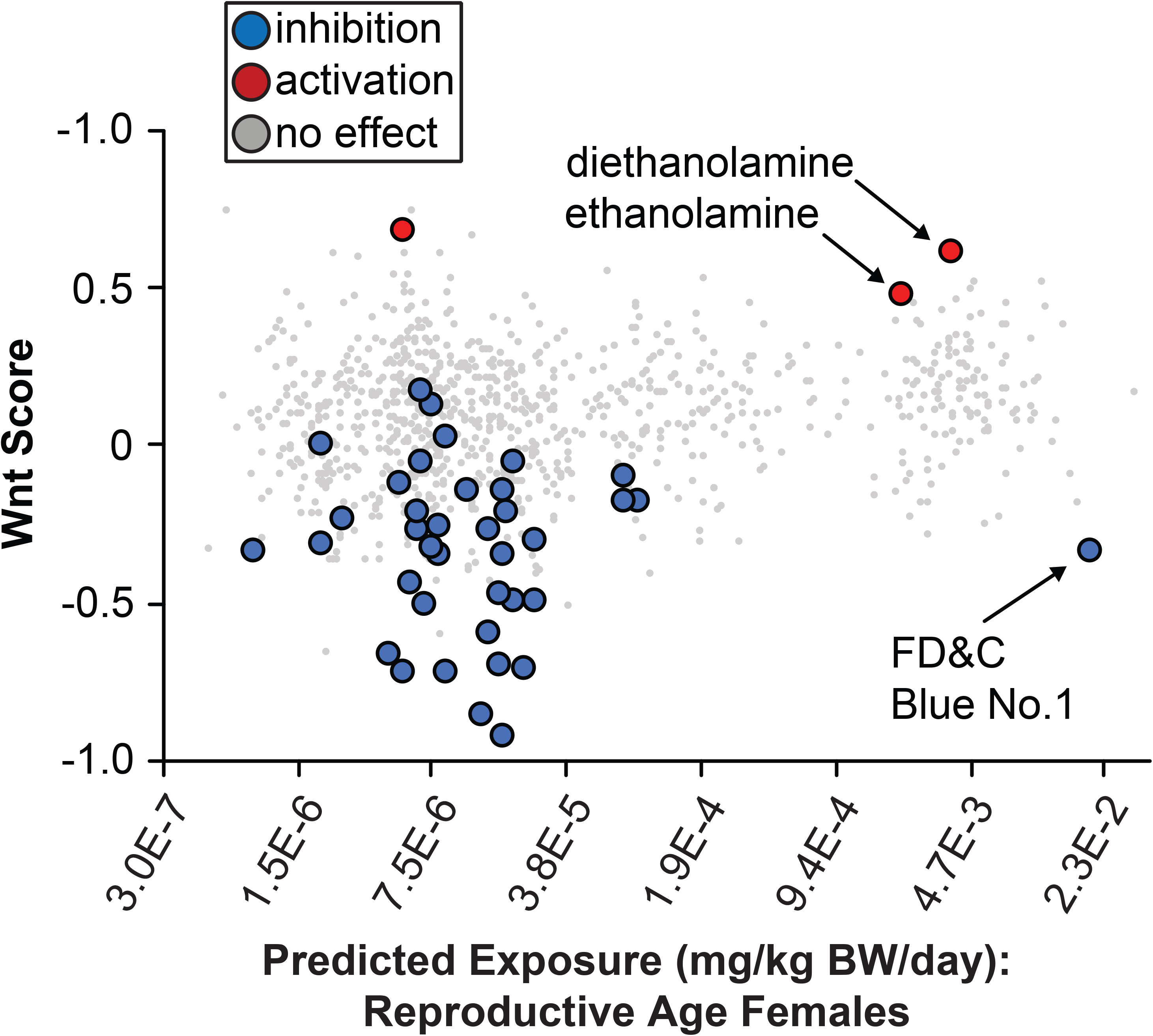
Human exposure prediction data. Predicted exposure of reproductive age females to ToxCast chemical libraries. Chemicals with non-toxic Wnt modulation from Figures 2,3 are colored.

The next Wnt modulator with high exposure predictions was DEA, which was in the 98th percentile of predicted exposure volume for all ~8,000 chemicals in the Tox21 set (Figure 4)(35). DEA is used in a wide range of products, including adhesives, printing inks, paint, pigments, and paper, among others (64). DEA is capable of absorbing through the skin, therefore the most likely route of human exposure is dermally through liquid laundry and dish detergents, shampoos, and soaps(64, 65), where it functions as a surfactant and pH adjuster(66). It is also used in manufacturing, where it is estimated that ~800,000 workers are exposed to DEA through occupations such as metalwork and road paving(64). There is inadequate epidemiological data for DEA exposure in humans, but DEA is classified as possibly carcinogenic in humans based on animal models(67), where dermal exposure demonstrates carcinogenic activity(68). DEA accumulates in specific tissues following repeat exposure, including the brain, where it is incorporated into phospholipids(69). DEA has also been shown to influence hippocampal neural progenitor proliferation at high doses *in vitro(70)* and *in vivo*(71, 72).

DEA is structurally similar to endogenous ethanolamine and choline. Cells and animals treated with DEA phenocopy choline deficiency, likely via competitive inhibition of choline metabolism(70, 73). However, there are no previous reports linking DEA to Wnt signaling, nor to any other developmental signaling pathways. For these reasons we decided to focus on DEA in follow up experiments. Using commercially obtained DEA, we tested the specificity of DEA in HEK293T cells against luciferase reporters that are sensitive to various signaling pathways. DEA concentration-dependently activated the Wnt reporter, with slight but statistically significant activation of the TGFβ reporter (Figure 5a). Wnt and TGFβ share many downstream target genes, and components of the two pathways are known to interact(74).

We next tested whether genetic background influenced the activity of DEA. We transfected HEK293T cells with either an empty plasmid, or one containing the autism linked *UBE3A^T485A^* mutant construct, and tested the effect of DEA on the Wnt reporter over a wide range of concentrations. Notably, DEA activated the Wnt reporter at 100-fold lower concentrations when transfected with *UBE3A^T485A^* (Figure 5b). DEA has previously been shown to decrease proliferation and increase apoptosis of mouse NPCs *in vitro* and *in vivo*(70, 71). *In vivo*, DEA affects hippocampal NPC proliferation at 80 mg/kg(71), which is substantially higher than what is predicted for human exposure (~0.0038 mg/kg bodyweight per day). Therefore, we sought to determine the lowest concentration at which DEA alters NPC proliferation using two genetically distinct phNPC lines(36). We compared DEA to known chemical Wnt modulators, including the Wnt activators CT99021, and lithium chloride. Each of the control chemicals increased proliferation as expected (Figure 5c,d). DEA increased proliferation in a concentration dependent fashion; the magnitude was similar to that of lithium chloride (Figure 5c,d). DEA was active at the lowest concentration tested (50 μM) in one cell line. We observed that higher concentrations were noticeably toxic to phNPCs (Figure 5d).

## Discussion

Here, we screened a library of environmental-use chemicals for their ability to modulate a Wnt sensitive reporter in cells overexpressing *UBE3A^T485A^*, an autism-linked gene that stimulates Wnt signaling at baseline. Previously, the EPA tested the ToxCast libraries for Wnt activation using a similar TCF7 reporter construct(75). Our approach is different for two reasons. First, in HEK293T cells TCF/LEF reporters are largely not expressed above baseline levels without additional treatment, which prevents detection of Wnt inhibitors. Second, we evaluated Wnt signaling in a genetically “sensitized” background, which we hypothesized would enhance the effects of Wnt modulators.

By comparing chemical structures, we identified classes of chemicals with shared effects on Wnt signaling, including synthetic estrogens, thyroid hormones, glucocorticoid and steroid hormones, and agricultural fungicides. Aside from PharmaGSID_48505, the primary cluster of non-toxic Wnt activators were ethanolamines, which are predicted to have relatively high levels of exposure in reproductive age females and children age 6-11 (Figure 4). We found that DEA specifically activated Wnt signaling in baseline conditions, but overexpressing the autism linked *UBE3A^T485A^* mutation amplified DEA’s effect on Wnt signaling (Figure 5). Consistent with the role on Wnt in regulating proliferation, we observed an increase in proliferation in primary human neural progenitor cells.

In animal models, DEA exposure has effects on several tissue/organ systems. Mice treated with DEA for two years develop higher rates of kidney and liver tumors (data reviewed in(65)). These tumors had high rates of mutations in exon two of the β-catenin gene, and demonstrated abnormal nuclear localization of β-catenin, indicative of constitutively active Wnt signaling(76). Topical treatment of DEA on pregnant mice reduces embryonic viability, and reduces proliferation of embryonic hippocampal neural progenitors *in vivo*(71). At high doses, DEA was found to reduce proliferation of cultured murine NPCs via inhibition of choline uptake(70). Choline is an essential nutrient crucial for normal brain development(77), and DEA affects patterns of DNA methylation that mimic choline deficiency(78).

There are several possible mechanisms by which DEA could modulate Wnt signaling. DEA is structurally similar to choline, and both are incorporated into phosphoglyceride and sphingomyelin analogs(69). Importantly, this unnatural incorporation includes a xenobiotic headgroup from DEA, and leads to bioaccumulation in the brain(69). The composition of the lipid membrane influences membrane localization and trafficking of lipid modified proteins(79), such as the Wnt ligands, which require lipid modification for proper solubility, the establishment of morphogen gradients, and protein interactions(80). Furthermore, the Wnt coreceptor LRP6 also uses phospholipids as signaling ligands(81). Thus, there are multiple possible mechanisms by which DEA exposure could influence lipid metabolism and affect Wnt signaling.

DEA is used in cosmetics due to its properties as a pH stabilizer (pH 9.5). Recent experiments in the context of chick development demonstrated that high intracellular pH caused by enhanced glycosylation leads to non-enzymatic ß-catenin acylation, which activates Wnt specific transcriptomic profiles that maintain mesoderm identity(82). Acidification of tumor cells also inhibits Wnt signaling in tumor cells(83). Furthermore, the loss of UBE3A affects pH of the Golgi apparatus, which compromises pH sensitive functions of the Golgi apparatus, such as glycosylation(84). Therefore, alterations in intracellular pH could be a mechanism by which DEA enhances Wnt signaling in the context of the hyperactive *UBE3A^T485A^* mutation.

The use of DEA in cosmetics was banned in Europe and Canada following concerns about DEA as a carcinogen(85, 86). The FDA and the National Toxicology Program have likewise found an association between DEA and cancer in lab animals, and provide information on the use of DEA and its derivatives in cosmetics (https://www.fda.gov/cosmetics/cosmetic-ingredients/diethanolamine). However, as of this writing, DEA is approved in the United States as long as it does not comprise >5% of the total product composition(65). To our knowledge there have been no epidemiological studies suggesting a role, or lack thereof, of DEA in increasing risk for neurodevelopmental disorders. Our data suggests that genetic background (i.e. *UBE3A^T485A^* expression) greatly enhances the effects of DEA on Wnt signaling (Figure 5b), and that DEA increases human NPC proliferation at relatively low concentrations (Figure 5c,d)(70). Given these findings, the predicted high level of exposure in humans, including women of childbearing age, additional studies are warranted, particularly with regard to exposure and neurodevelopmental outcomes in genetically sensitized backgrounds.

## Acknowledgements

We thank Tammy Havener and the UNC Catalyst for Rare diseases for use of the high-throughput screening facility, Thomas Girke for technical assistance, John Wambaugh for providing human exposure prediction data, and Steven Zeisel for his helpful comments.

## Contributions

JMW and MJZ designed the experiments and wrote the manuscript. JMW performed all experiments and analysis. JJ performed the experiments in Figure 5A. JLS provided reagents and protocols for performing human neural progenitor cultures.

## Figure Legends

**Supporting Figure 1.**
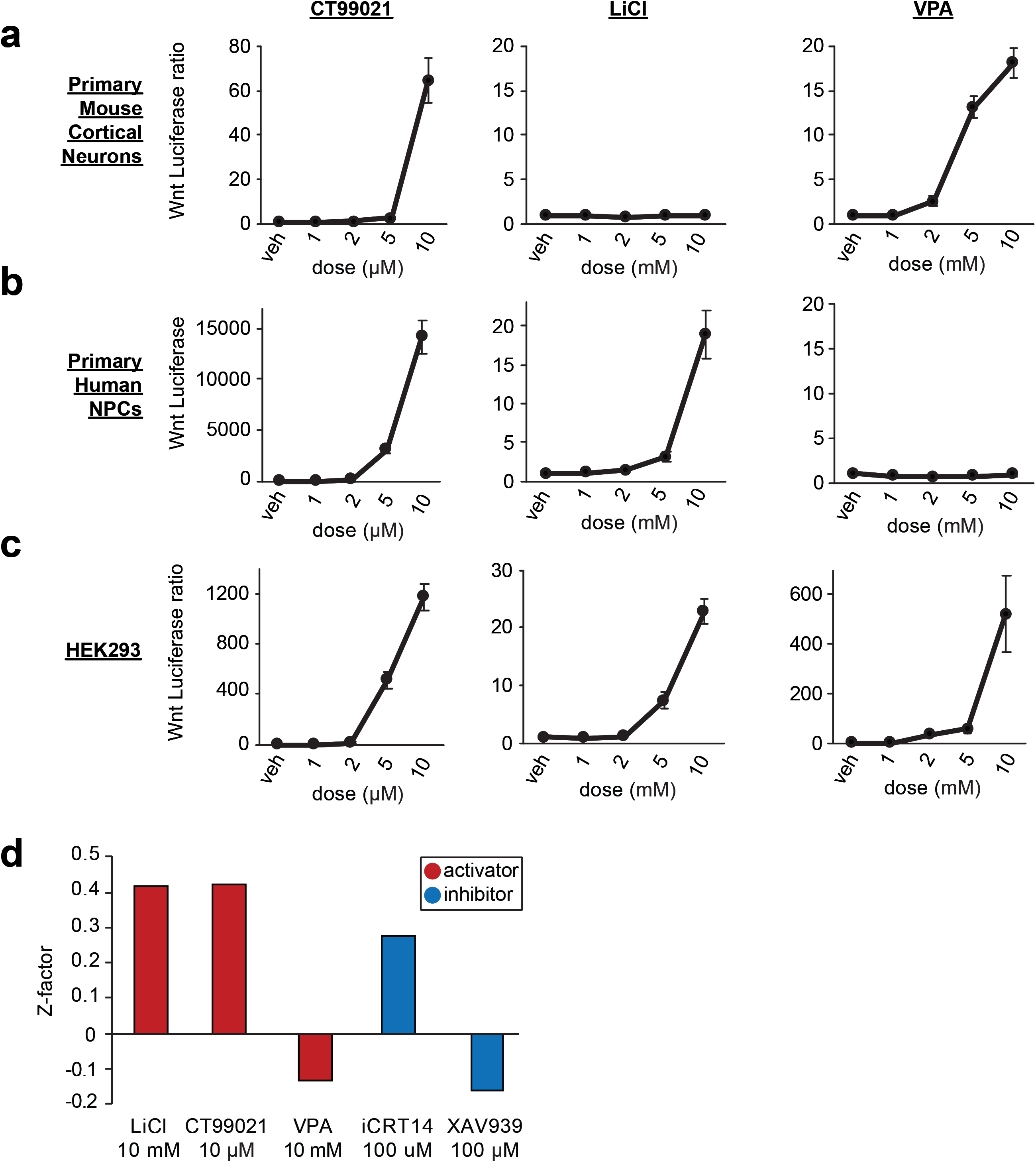
Establishing ToxCast screen conditions. **a-c)** Chemicals known to activate Wnt signaling tested in primary mouse cortical neurons (**a**), primary human neural progenitor cells (**b**), and HEK293T cells (**c**), without *UBE3A^T485A^* overexpression. Primary cells were transduced with lentiviruses carrying BAR:Firefly and Tk:*Renilla*. HEK293T cells were transiently transfected with plasmids. n=4 per condition. **d)** Z-factor analysis for known Wnt activating and inhibiting control chemicals at indicated concentrations.

**Supporting Figure 2.**
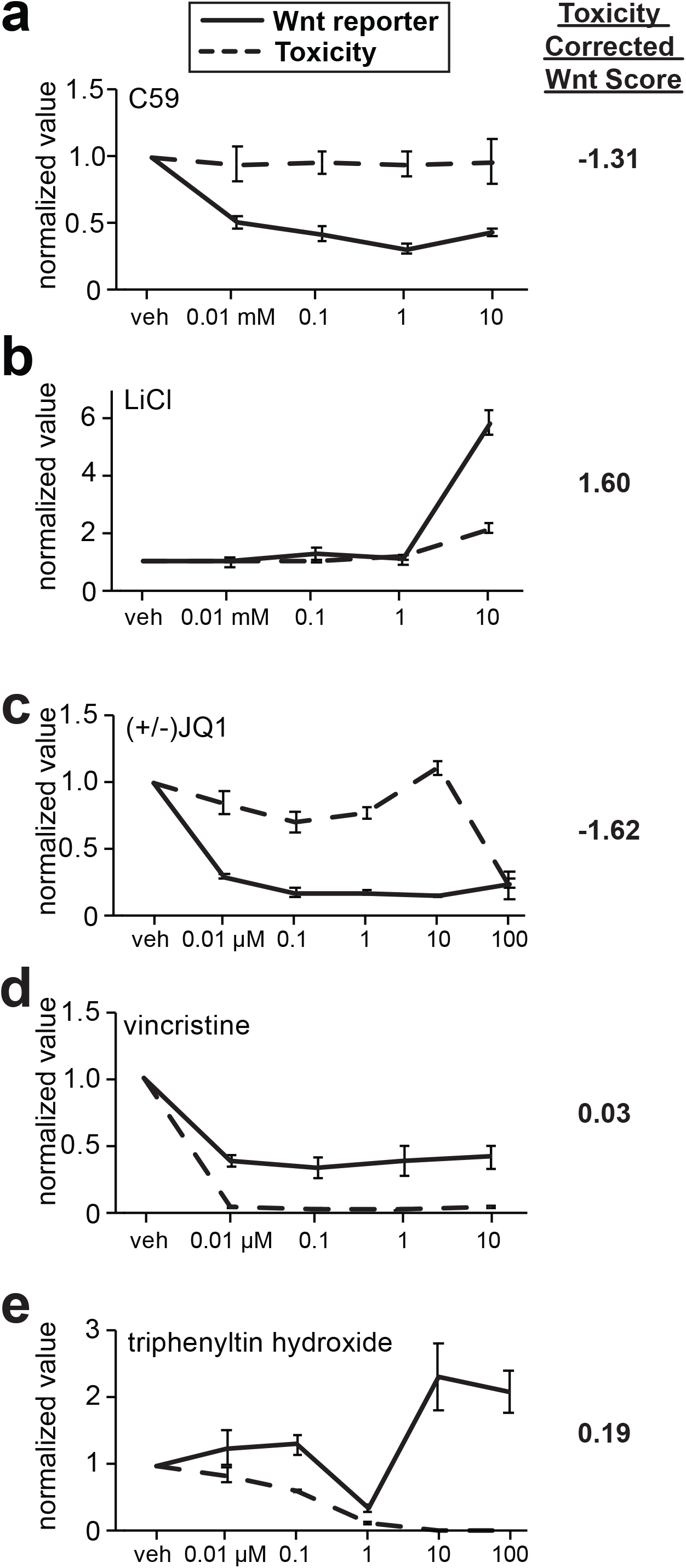
Examples of concentration-response curves and Wnt Score. **a-c)** Control chemicals that modulate Wnt reporter (solid line) at concentrations that are non-toxic (dashed line). The Wnt score listed to the right of each graph is a single score combining multiple concentrations of the Wnt reporter values with a penalty for toxicity (see methods). **d,e)** Concentration-response curves for two ToxCast chemicals that significantly inhibit (**d**) or activate (**e**) the Wnt reporter, but do so at concentrations that are toxic. The Wnt score listed to the right, centered around 0, reflects the penalty incurred on Wnt activation from toxicity.

**Supporting Figure 3.**
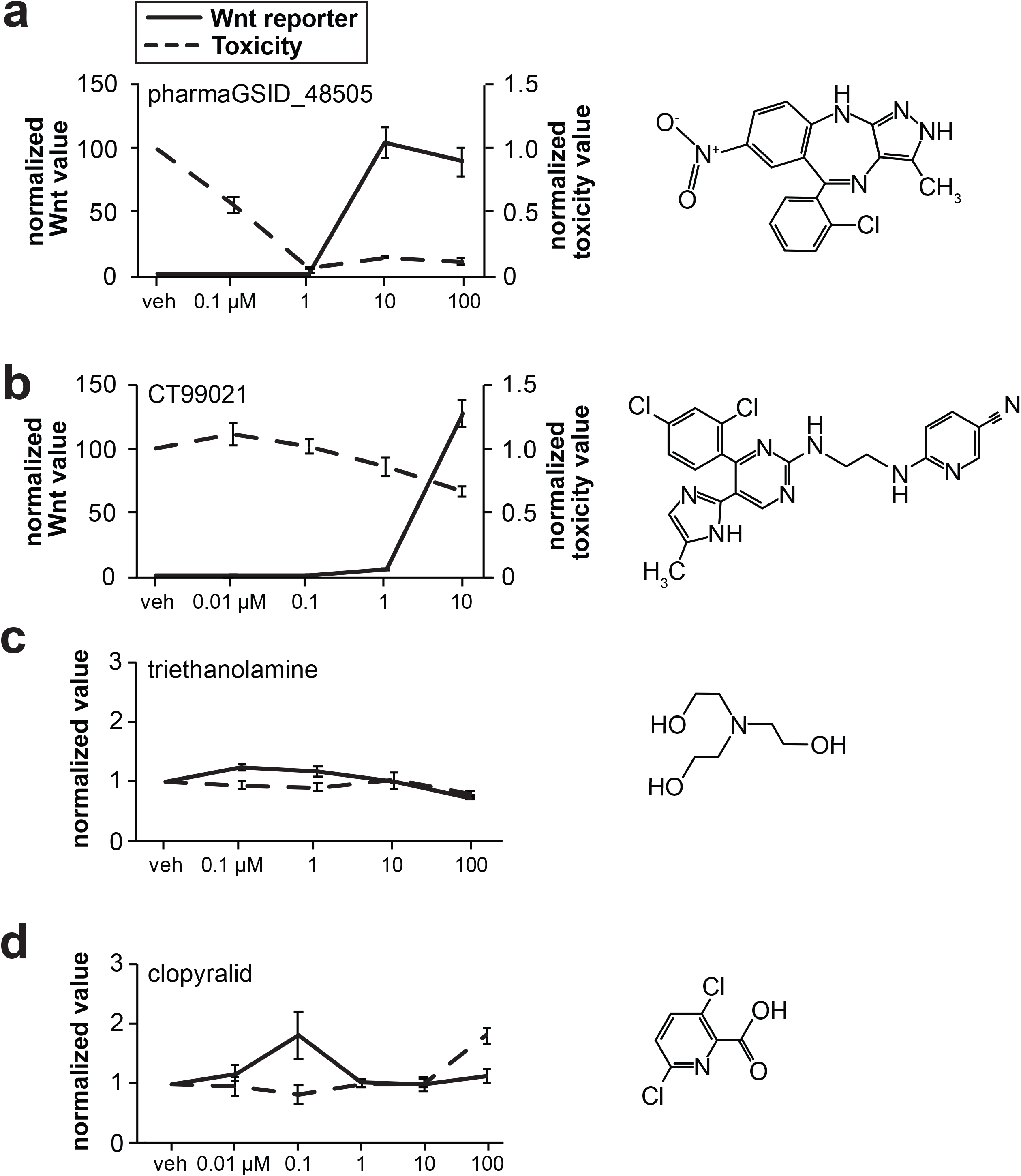
Concentration-response curves for two Wnt activating chemicals and two chemicals that are structurally similar to activators, but that fail to activate Wnt signaling. **a-d)** HEK293T cells transfected with *UBE3A^T485A^* treated with **(a)** the most potent ToxCast wnt activator pharmGSID_48505, and **(b)** CT99021, the most potent Wnt activator identified to date. Results demonstrate similar effect sizes with higher toxicity for pharmaGSID_48505. **(c)** Triethanolamine, the trimeric form of monoethanolamine, fails to activate Wnt reporter. **(d)** Clopyralid alone also fails to activate Wnt reporter.

## Notes

### Competing Interest Statement

The authors have declared no competing interest.

